# Harnessing formal concepts of biological mechanism to analyze human disease

**DOI:** 10.1101/350371

**Authors:** Lindley Darden, Kunal Kundu, Lipika R. Pal, John Moult

## Abstract

Mechanism is a widely used concept in biology: in 2017 more than 10% of PubMed abstracts used the term. Thus, searching for and reasoning about mechanisms is fundamental to much of biomedical research, but until now there has been almost no computational infrastructure for this purpose. Recent work in the philosophy of science has explored the central role that the search for mechanistic accounts of biological phenomena plays in biomedical research, providing a conceptual basis for representing and analyzing biological mechanism. The foundational categories for components of mechanisms - entities and activities - guide the development of general, abstract types of biological mechanism parts. Building on that analysis, we have developed a formal framework for describing and representing biological mechanism, MecCog, and applied it to describing mechanisms underlying human genetic disease. Mechanisms are depicted using a graphical notation. Key features are assignment of mechanism components to stages of biological organization and classes; visual representation of uncertainty, ignorance, and ambiguity; and tight integration with literature sources. The MecCog framework facilitates analysis of many aspects of disease mechanism, including the prioritization of future experiments, probing of gene-drug and gene-environment interactions, identification of possible new drug targets, personalized drug choice, analysis of non-linear interactions between relevant genetic loci, and classification of disease based on mechanism.

## INTRODUCTION

In the pre-database era, data were scattered throughout the literature, and many projects were hampered by the difficulties of discovering and tabulating data. Databases have transformed our ability to retrieve and work with this class of information. A similar, more severe problem now exists for the retrieval and comprehension of biological facts: Over a million papers on biology are published each year, each recording details about biological systems. How can we extract, organize, and represent this knowledge to maximize its accessibility and utilization? We address this problem using formal concepts of mechanism that have emerged from the philosophy of biology [1].

Philosophers have characterized mechanisms in terms of entities and activities organized to produce a phenomenon [1]. The central insight from that work is that fundamentally, all mechanisms are composed of activities between two or more entities. In biological systems, these entities may be molecules, or cells, or things at other levels of biological organization. The activities result in altered entities. For example, an amino acid substitution in a protein entity may result in an activity of a weaker intra-molecular interaction, so perturbing the protein’s abundance; a protease entity may have the activity of cleaving the polypeptide backbone of another protein entity, so altering it; a cell entity may emit a cytokine which binds to a receptor on another cell entity, so altering the latter’s transcriptional profile. In the MecCog framework, altered entities are termed “substate perturbations” (SSPs) and activities or groups of entities and activities, are termed “mechanism modules” (MMs). Biological mechanisms may be described by a series of steps in which at each step the activity of an entity alters the state of another entity. A key feature is the inclusion of specific activities that drive the changes from step to step. This viewpoint - asking at each step of a mechanism what are the entities and activities - guides and constrains the search for a mechanism’s salient features.

In contrast to biological pathway representations (for example KEGG [2], Reactome [3], WikiPathways [4]) which describe relationships between system components, MecCog is built around the productive mechanism modules that drive system changes. MecCog focuses on the perturbations of biological systems that lead to a disease outcome, rather than the extensive systems’ description used in Aetionomy [5], the Parkinson’s Disease Map [6], and the SBML pathway project Payao [7]. MecCog makes a clear distinction between mechanism modules and biological entities, in contrast to other emerging mechanism-oriented representation projects such as Noctua (http://noctua.berkeleybop.org/). MecCog’s causal chain related approach is shared by the Collaborative Adverse Outcome Pathway Wiki (AOP-Wiki) [8] initiative, a crowdsourced representation of toxicology pathways that integrates perturbed entity information from molecular to cellular to organ biological scales, although AOP does not explicitly represent mechanism. MecCog extends representation of evidence for mechanism components such as that provided by the Evidence and Conclusion Ontology (ECO) [9] to include levels of confidence in mechanism components, ambiguities in mechanism schemas, and different types of ignorance.

MecCog focuses on the relationship between genetic variation and disease phenotypes, but also includes mechanisms arising from a drug intervention or an environmental change. MecCog can be used to represent all types of human disease with a genetic component, including rare monogenic disease, cancer, and common complex trait disease. In most respects, the requirements for building a mechanism schema are the same across these disease types, and so only a single disease mechanism framework is required. In common with most pathway and disease mechanism representations, MecCog schemas are compiled manually. To this end, the infrastructure has been designed to maximize ease of input and evidence recording.

## REPRESENTATION OF DISEASE MECHANISM

Each disease mechanism schema begins with a genetic variant, a drug intervention, or an environmental change. Here we focus on genetic perturbations. For monogenic diseases, mutations that have been shown to be causative (i.e., are ‘pathogenic’); for cancer, driver mutations; and for complex trait disease, SNPs associated with a disease phenotype, typically derived from a Genome Wide Association Study (GWAS) [10].

For all types of disease, each such situation signals the presence of a biological mechanism linking the perturbation at the DNA stage (an altered base for example) to a perturbation at the disease phenotype stage (for example altered disease risk) (Figure 1).

**Figure 1.**
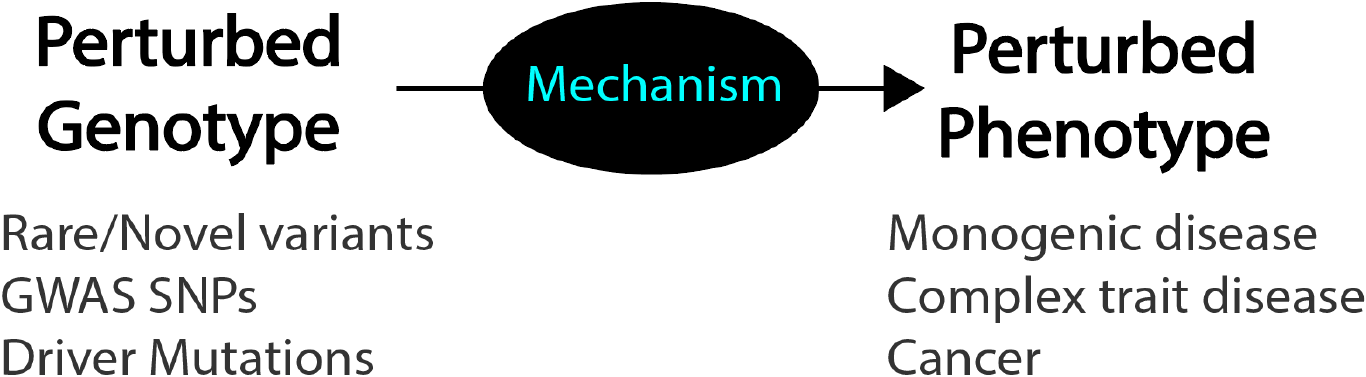
Every relationship between the presence of a genetic perturbation and a disease phenotype implies the presence of an underlying mechanism. Without further information, that mechanism is unknown – a ‘black box’.

Multiple mechanism steps link a genetic variant and a disease phenotype. Each step in the mechanism schema consists of a triplet of an input substate perturbation, a mechanism module, and an output substate perturbation (SSP-MM-SSP) (Figure 2), a special case of a subject-predicate-object triplet, where the predicate is an active verb. The output SSP of a step forms the input SSP to the next step. Possible and existing sites of drug intervention are depicted by octagons, and environmental changes by a cloud symbol.

**Figure 2.**
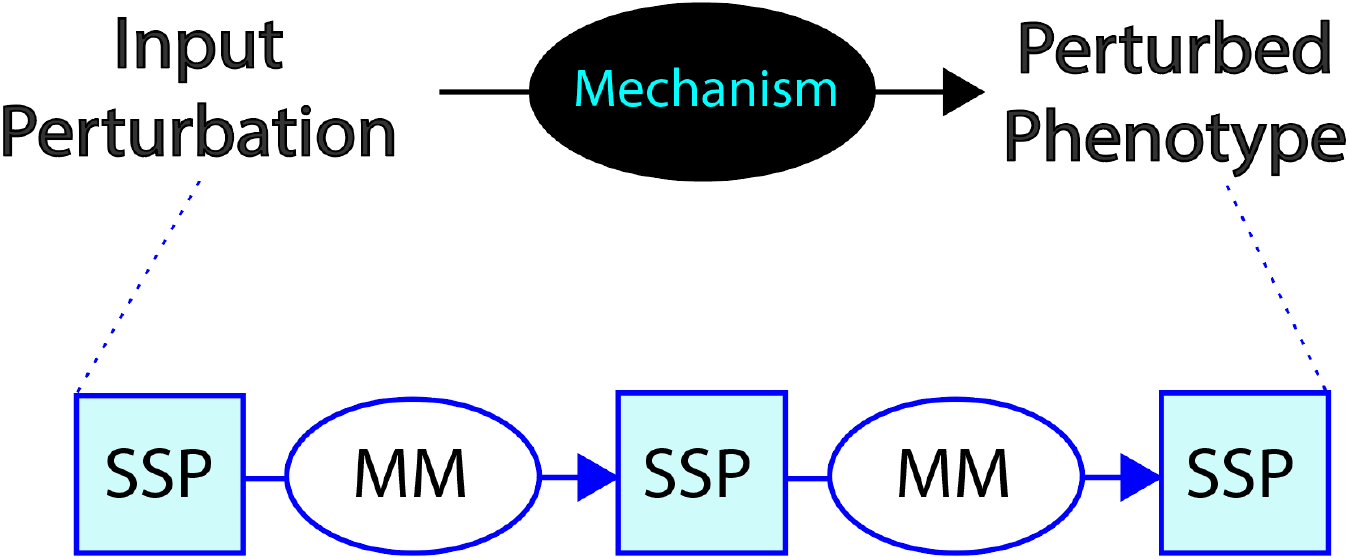
A mechanism schema consists of a series of steps. Each step has an input substate perturbation (SSP), a mechanism module (MM), and an output substate perturbation.

### Schema steps and stages

The mechanism module in a step describes the activity or mechanism (composed of entities and activities) by which the input substate perturbation produces the output substate perturbation. For example, an amino acid substitution in a protein (a substate perturbation) may result in lower abundance of protein complex (a resulting substrate perturbation) through the mechanism module of weaker interactions (the activity) between the two proteins (the entities). Each perturbation occurs at one of a small number of organizational stages: DNA, RNA, Protein, Macromolecular complex, Organelle, Cell, Tissue, Organ, or Organism phenotype. For example, a DNA variant may result in a lower transcription rate for a gene resulting in less messenger RNA, in turn resulting in less protein product, and as a consequence less of a protein complex involved in signaling the presence of bacteria, resulting in an altered immune cell response, and so on. There may be more than one step in a given stage (for example, multiple protein perturbation steps) or no steps at a specific stage (for example, an RNA molecule directly affects a cell state, without the involvement of a protein). The schema formalism permits telescoping of sets of steps into a single mechanism module, so that well-established sub-mechanisms need not be spelled out in detail, for example ‘protein synthesis’, again increasing focus on the disease-related perturbations.

### Mechanism component classes

At each stage there is a small number of likely classes of perturbation. For example, at the DNA stage, there are four perturbation classes: single nucleotide variant (SNV), insertion or deletion (IN/DEL), copy number variation (CNV), and chromosomal rearrangement. Similarly, within a stage, or between a pair of stages, there are a small number of likely mechanism module classes. For example, a perturbation at the RNA stage may lead to the next perturbation through one of five mechanism modules: Altered intra-RNA interactions, altered RNA/RNA interactions, altered RNA/protein interactions, altered RNA/splicing factor interactions, and altered RNA editing. Where possible, class names are taken from existing ontologies, currently GO [11], Sequence Ontology [12], Variation Ontology [13], NCIT [14], MESH [15], with the addition of the modifiers ‘increased’, ‘decreased, ‘no’, ‘altered’, ‘sooner’, ‘later’. The delineation of a set of organization stages in a mechanism and the classes of possible mechanism components at each stage is a powerful aid to schema construction.

### Representation of ignorance, uncertainty, and evidence

A key feature is the representation of ignorance, alternatives, ambiguities, and uncertainty, linked to supporting evidence (Figure 3). Unknown mechanisms linking two substate perturbations are depicted as black mechanism modules. Where there is uncertainty as to whether such a link exists, the black mechanism module contains a question mark. Ambiguities or alternatives in a schema are shown as AND, OR, or AND/OR branches, and the level of uncertainty is color coded.

**Figure 3.**
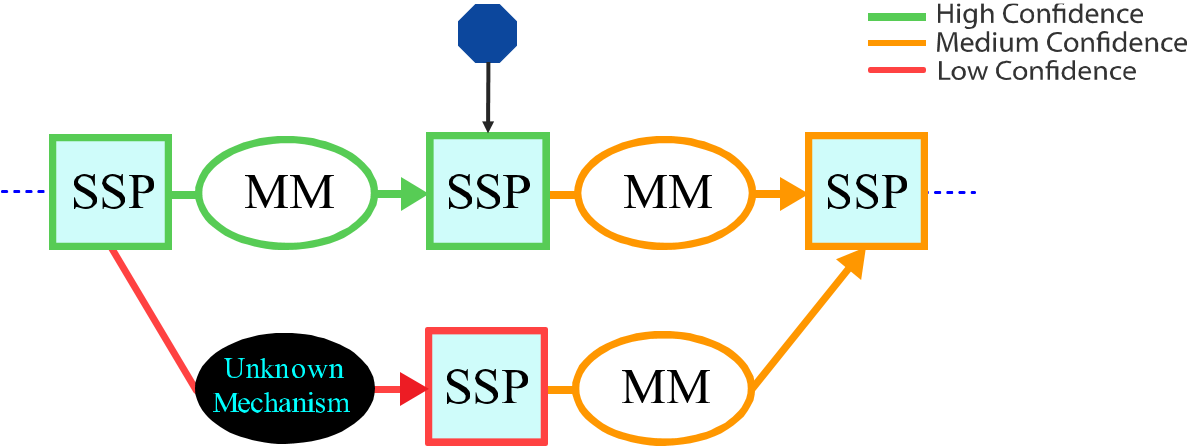
Representation of Ignorance, ambiguity, and uncertainty. Unknown mechanism components are represented by black ovals; ambiguity by branching in the schema; and uncertainty of evidence by element color (green for high confidence, orange for medium confidence, and red for low confidence). Blue octagons represent sites of possible therapeutic intervention.

For each symbol, popup boxes give access to more detailed information, and link to fuller entries that provide a brief commentary on the mechanism feature, a summary of the evidence and the evidence sources, links to the relevant literature and data, and a confidence value (1 – 5). These conventions are consistent with but extend beyond the emerging confidence information ontology [16], particularly in the use of branching within the schema to represent ‘and/or’ uncertainty, and the explicit representation of mechanism ignorance.

### Disease mechanism graphs

For complex trait disease and for cancer, multiple genetic perturbations at multiple loci contribute to a disease, and each such relationship is represented by a mechanism schema. For a particular disease, there are also schemas representing how each drug affects disease phenotypes. There may also be schemas for environmental effects. For example, in Crohn’s disease, there are contributions to disease risk and other disease-related phenotypes from variations in microbiome composition [17]. Schemas for different loci may have substate perturbations in common, for example, the ‘innate immune response’ SSP in the MSP locus schema (see below) is shared by multiple other loci, such as NOD2 [18] and ATG16L1 [19]. Thus, the full set of schemas for a disease form a disease mechanism graph. As discussed below, these graphs enable a number of applications.

### Expert sourcing

Eventually, we anticipate that mechanism schemas will be populated directly from mining the literature, by identification of subject-object-predicate triplets corresponding to the SSP-MM-SSP triplets of schemas. The current state of the art for this kind of text mining, as reflected by objective testing in the BioCreative community challenges [20], is such that this is not yet practical. Thus we have optimized the MecCog resource for human interaction, with an intuitive graphical language, and extensive tools to facilitate construction, including pull-down menus of possible classes for each mechanism stage.

## COMPUTATIONAL INFRASTRUCTURE

MecCog is implemented on Node.js, a JavaScript runtime built on Chrome’s V8 JavaScript engine. Node.js uses an event-driven non-blocking I/O model and so is lightweight and efficient. The web application is built using Sails.js, which provides a web framework to build custom enterprise-grade Node.js applications. Sails also has an Object-relational mapping (ORM), Waterline, providing a simple data access layer for different types of backend database. The MecCog implementation uses the MySQL relational database management system to store data on users and mechanism schemas. All the database transactions use REST APIs and are secured by the CSRF (Cross-site request forgery) protection feature in Sails. The interface to build, edit and view mechanism schemas is powered by the Rappid Diagramming Framework (https://www.jointjs.com/), written in JavaScript. Rappid’s feature of communicating with databases via AJAX and JSON makes it compatible with the database access layer provided by Waterline in Sails.js. The aesthetics of the website is supported by the open-source front-end library Bootstrap.js. As the project progresses, all mechanism schemas will be encoded by the network modeling language - Biological Expression Language (BEL - http://openbel.org) in order to represent the schemas in a computable form.

## EXAMPLE MECHANISM SCHEMA FOR A COMPLEX TRAIT DISEASE LOCUS

As noted earlier, the mechanism schema framework can be used to describe and analyze the relationship between genetic variation and disease phenotypes for all types of genetic disease. Examples are available on the MecCog website (www.meccoq.org). Here we describe one case of a locus implicated in risk of a complex trait disease. GWA studies have now revealed over 160 loci scattered throughout the genome where the presence of a SNP (single nucleotide variant (a SNP) is associated with increased risk of Crohn’s disease [21]. For some loci, the corresponding mechanisms have been extensively studied, for example [18, 19]. For others, little or nothing is yet known. Figure 4 shows an example of a mechanism schema for a moderately well understood Crohn’s disease locus - the relationship between the presence of a GWAS marker SNP (rs3197999) associated with increased risk of the disease [22] via a mechanism affecting the activity of Macrophage Stimulating Protein (MSP). The mechanism begins at the perturbed DNA substate (the presence of a missense SNP), and then protein, protein-protein complex, cell signaling, inflammatory response, and the gut wall steps, to the final effect on disease risk.

**Figure 4.**
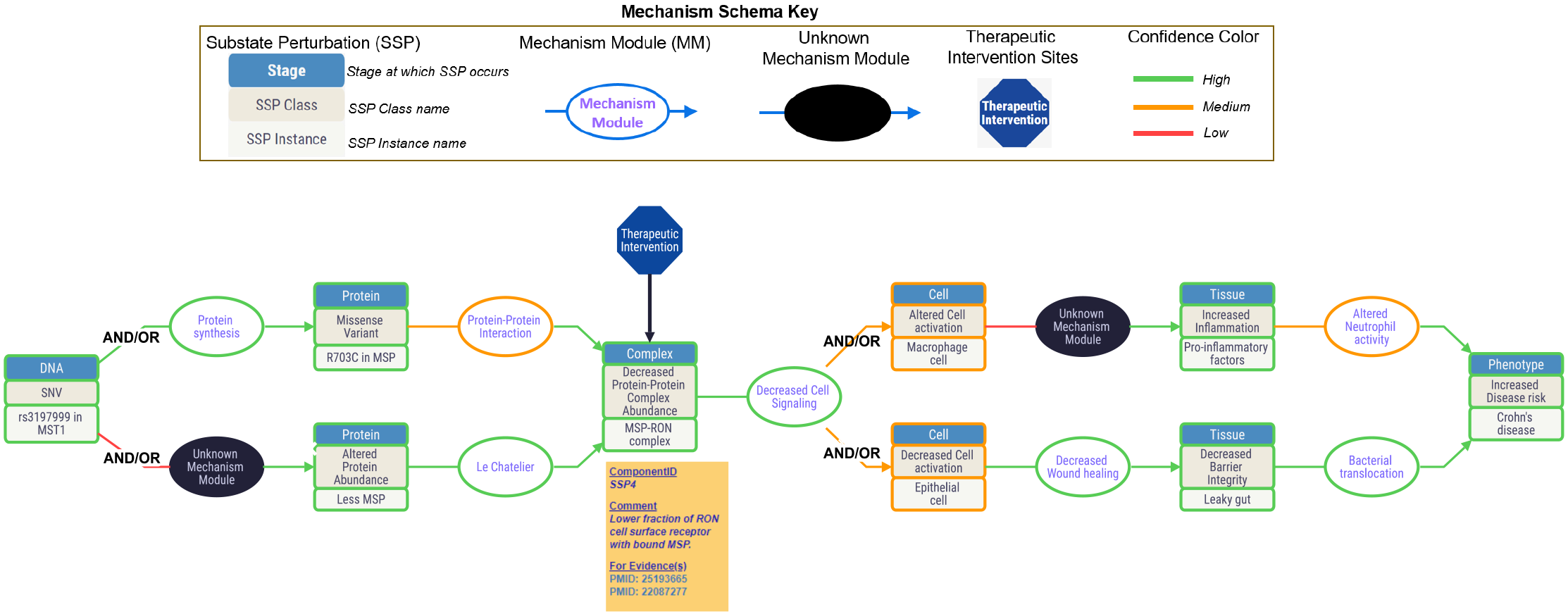
A disease mechanism schema, relating the presence of a missense SNP in Macrophage Stimulating Protein (MSP) to increased risk of Crohn’s disease.

This schema contains two unknown mechanism modules (black ovals). For one of these, experiment has shown a lower serum abundance of the MSP in the presence of the disease risk marker SNP, but the mechanism for that is unknown (is it decreased expression, lower protein stability, altered degradation properties, for instance). In the other, altered macrophage activation results in an increased inflammatory response, but the exact mechanism is unclear. Confidence in the schema steps varies: high (green), medium (orange), and low (red). There are also two major mechanism ambiguities, represented by the two branched sections of the schema. The first branching reflects the facts that one study [23] has reported that the presence of mechanism SNP results in a weaker protein-protein interaction between MSP and a cell surface receptor, RON, while another study [24] reports that the SNP affects the level of MSP protein in serum but not the binding affinity between MSP and RON. The two branches lead to the same outcome – a lower concentration of the MSP-RON protein complex. The second branch in the schema (the parallel paths in the right-hand segment) represents uncertainty as to whether the lower intra-cellular signaling resulting from reduced abundance of the MSP-RON complex most affects macrophage activity and consequently innate immunity; or whether it affects wound healing activities of epithelial cells, so primarily alters barrier integrity; or both.

Many schemas, for example that for variants in NOD2 [18] (accessible on the MecCog site), are substantially more complicated. All schemas examined exhibit knowledge gaps and ambiguities.

## APPLICATIONS OF DISEASE MECHANISM SCHEMAS

In addition to providing an integrative framework for describing what is and is not known about a disease mechanism, the schemas have a number of further applications:

### Prioritizing future experiments

Identification of gaps and uncertainty in knowledge allows a more objective evaluation of which experiments are most critical. For example, of the two major ambiguities in the MSP schema, the first - whether the mechanism SNP results in less MSP protein or a weaker MSP-RON complex, or both - has no major implications for the disease mechanism because subsequent steps in the schema are unaffected. But the second ambiguity - whether it is innate immune cell activation or wound healing that is affected, or both - is important for understanding how this risk factor fits into the overall picture of the disease, and suggests that further experiments to resolve the ambiguity are warranted.

### Identifying possible new sites of therapeutic intervention

Schemas also allow identification of potential sites of therapeutic intervention. For example, in spite of the uncertainties and ambiguities in the MSP schema, the central role of the MSP-RON complex is clear, and that suggests a possible therapeutic intervention: an appropriate compound (a conventional small molecule drug, or an antibody) that bridges the structural interface between RON and MSP [23] could restore wild-type signaling strength. Of course there are many reasons why this may turn out not to be a useful target; but the mechanism schemas do provide a means of systematically identifying such possibilities.

### Epistatic (non-additive) effects

Non-additive contributions from pairs or higher order combinations of variants contributing to complex trait disease are widely expected to play a major role in disease mechanisms [25]. In cancer, identification of such interactions has provided the basis of a treatment strategy [26], and a similar strategy may be possible for complex trait disease, if the interactions can be found. In principle, GWAS data can be analyzed to discover nonadditive relationships between genetic variants. In practice, the large number of possible combinations drowns any potential signal (a study looking at N variants implies N^2^ statistical tests), and very large studies will be needed to overcome that. The mechanism graph for a disease, formed by combining all relevant schemas, facilitates the generation of specific hypotheses that can be tested against GWAS data with minimal multi-testing issues. For example, in Crohn’s disease, the combined graph for loci relevant to bacterial penetration of the gut lining mucosal layer emphasizes a number of possible non-linear effects between mucin gene variants affecting mucosal layer integrity [27] (MUC1, MUC2), variants affecting the unfolded protein response [28] (XBP1, ORMDL3), and variants affecting autophagy [29] (NOD2, ATG16L1, LRRK2, IRGM).

### Precision Medicine

For complex trait diseases and for cancer, each affected individual has contributing variants that affect only a subset of disease mechanism related genes. For instance, an individual diagnosed with Crohn’s disease typically has risk alleles in approximately half of the known risk loci. Thus, for each patient, only part of the full disease mechanism graph is relevant, and the extent and nature of the overlap of that subgraph with the schemas for available drugs provides a potential means of prioritizing drug choice. For example, an antibody inhibitor of SMAD7, Mongersen, is an effective treatment for about 65% of Crohn’s patients [30]. Mongersen acts by modulating a negative feedback loop by which SMAD7 reduces the anti-inflammatory response to TGFβ1 [30]. The mechanism schema for this drug links the likely effectiveness of Mongersen to Crohn’s marker SNPs in the Toll Like Receptor 4 (TLR4) and αVβ8 integrin loci, as well as the more obvious relationship to the SMAD7 marker. In principle, association studies of the relationship between drug efficacy and marker SNPs can also provide this information, for instance [31], but because of the large number of possible associations that must be tested, there are often insufficient data to provide robust associations (for example [32]). Restricting tests of statistical significance to drug/variant hypotheses generated by schema analysis will mitigate multi-testing correction effects, and so increase the power of the association studies.

### Disease classification

Historically, diseases have been classified on a number of criteria: the location of the disease (anatomical), the cause of the disease (etiology), or the symptoms of the disease [33]. The disease ontology [34] utilizes these criteria, as well as others. It has also been proposed that a disease taxonomy should be based on molecular mechanisms [35], so that patient subgroups can be characterized by their shared molecular etiology. The set of classes for each stage of mechanism schemas provides a basis for a mechanism-based comparison of genetic diseases. For example, which disease mechanisms involve lower abundance of a protein? Where that occurs, is it produced by lower expression, shorter half-life, or impaired folding? Where the latter is the case, is it mediated by altered ubiquitination rate? Which disease mechanisms include altered protein-protein interactions, which include altered cell-cell signaling, and so on. In addition to providing increased insight into the mechanistic basis of disease, this classification may also help delineate therapeutic strategies that may be applicable in each case.

## ACKNOWLEDGEMENTS

This work was supported in part by NIH R01GM104436 to JM. We thank Yizhou Yin for many key comments and suggestions.

